# Automatic fruit morphology phenome and genetic analysis: An application in the octoploid strawberry

**DOI:** 10.1101/2020.11.09.374744

**Authors:** L.M. Zingaretti, A. Monfort, M. Pérez-Enciso

## Abstract

Automatizing phenotype measurement is needed to increase plant breeding efficiency. Morphological traits are relevant in many fruit breeding programs, as appearance influences consumer preference. Often, these traits are manually or semi-automatically obtained. Yet, fruit morphology evaluation can be boosted by resorting to fully automatized procedures and digital images provide a cost-effective opportunity for this purpose. Here, we present an automatized pipeline for comprehensive phenomic and genetic analysis of morphology traits extracted from internal and external strawberry images. The pipeline segments, classifies and labels the images, extracts conformation features, including linear (area, perimeter, height, width, circularity, shape descriptor, ratio between height and width) and multivariate (Fourier Elliptical components and Generalized Procrustes) statistics. Internal color patterns are obtained using an autoencoder to smooth out the image. In addition, we develop a variational autoencoder to automatically detect the most likely number of underlying shapes. Bayesian modeling is employed to estimate both additive and dominant effects for all traits. As expected, conformational traits are clearly heritable. Interestingly, dominance variance is higher than the additive component for most of the traits. Overall, we show that fruit shape and color can be quickly and automatically evaluated and is moderately heritable. Although we study the strawberry species, the algorithm can be applied to other fruits, as shown in the GitHub repository https://github.com/lauzingaretti/DeepAFS.

## Introduction

Demographic pressure and climate change are two of the major challenges of the 21st century. The worldwide population continues growing exponentially and it is expected to reach ∼9.8 x 10^9^ in 2050 [1]. Climate change generated by greenhouse gas emissions is possibly the greatest threat, as it is leading to extreme weather conditions, increasing areas of drought, and species extinction, among others [2–4]. In this adverse context, food production needs to be increased significantly. Increasing food production is not enough though, food safety and environmental care need to be preserved as well, i.e., a novel approach in breeding programs is required [5,6].

Artificial breeding has been a main responsible for the dramatic rise in food production that we have witnessed for over a century. The main goal of plant and animal breeding is to utilize genetic variability of complex traits to increase performance and optimize use of resources. A current bottleneck in plant breeding programs is the evaluation of hundreds of lines under different environmental conditions [7,8]. Plant breeding involves both genomics and phenomics, i.e., the expression of a genome in given environments. While available technologies can routinely and inexpensively scan the genome, characterizing high-throughput phenotypes remains a difficult task [9,10]. Automatizing phenotype measurement is then needed to increase the pace of artificial selection and is, unsurprisingly, one of the main targets of ‘Precision Agriculture’ [11,12].

The term ‘phenomics’ or ‘phenometrics’ was coined by Schork [13] as an attempt to understand events happening in between full genome and clinical endpoints phenotypes in complex human diseases. The expression quickly spread to animal and plant breeding research as a concept that bridges the gap between genotypes and the ‘end-phenotypes’. Although the term phenomics was devised in line with ‘genomics’, that is, to describe the whole phenome of any organism, note the phenome varies over time and between cells or tissues, and can never be fully portrayed [14].

A window of opportunities has emerged in the phenomics field with the recent development of robotics, electronics, and computer science. The subjective, time-consuming and often destructive human data collection is being replaced by sensors, digital cameras, cell phones, unmanned aerial vehicles (UAV), mass spectrometry, among others, that allow collecting hundreds of phenotype data objectively and inexpensively [9,15–17]. The challenge now is to develop new and improved analytical tools, capable of guaranteeing the transformation of all this wealth of data into valuable knowledge [15].

Digital images are among the cheapest and most widely available type of data. Imaging allows assessing morphological traits, which are highly relevant in numerous plant breeding schemes, since they can critically affect consumer acceptance especially in fruits [18–20]. Nevertheless, consumer preferences on appearance traits differ between communities around the world. Like most traits, fruit shape is determined by genetic and environmental factors such as flower morphology or insect-mediated pollination [21,22]. In all, morphological traits are among those with the highest heritability, which has allowed breeders to rapidly modify shape, size, and color patterns of agricultural products.

Although numerous works have been developed in the area of fruit morphology [23–26], most of them have focused in the inheritance of linear measures, e.g., diameter, perimeter, circularity, etc. By definition, however, morphological traits are highly dimensional. Computing only linear, univariate phenotype leads to a loss of information by extremely simplifying the features of a shape [27,28]. The use of geometric-morphometric approaches for shape analysis is warranted [29].

Fruit shape has been traditionally evaluated subjectively [30], but it can be boosted by resorting to automatized procedures. For instance, obtaining hundreds of fruit pictures can be routinely and inexpensively collected, even in the field, with a cell phone camera. Automatized image processing and analysis can then dramatically change the way shape and color traits are collected and characterized.

Here, we present a comprehensive phenomic and genetic analysis pipeline for fruit morphology automatic analysis. Two main issues are addressed: 1) converting the raw data (fruit images) into a processed curated database, and 2) designing an efficient analysis workflow to analyze fruit shape phenome. Finally, genetic parameters can be automatically inferred, either from pedigree or marker information. We apply the pipeline to images of cultivated strawberry (*Fragaria x ananassa*) fruits. Shape varied between the cultivars studied, e.g., circular, ellipsoid or rhomboid and color ranged from white to dark red. Our workflow establishes a proof of concept, which can be easily transferred to other visual phenotypes and fruits with minor modifications.

## Materials and Methods

### Plant material and Imaging acquisition

Strawberries were grown in plastic semi-tunnel using standard cultivation practices in South West Spain (Huelva, 37° 16’ 59’’ N, 7° 9’ 18’’ W) by PLANASA breeding company (https://planasa.com/en/). The experiment consisted of 24 crosses between 30 parental lines of F. x ananassa, for which 20 randomly chosen lines per cross were evaluated, giving a total of 508 genotypes. Fruits were collected from two individual plants of each line at the end of April 2018 in only one harvest event. We took images of 1 to 7 sliced fruits per genotype using a Nikon D80 digital camera. Samples were laid on a black surface, with the camera positioned at 35 cm height. The focal length was 18 mm, the manual aperture was f/8, and the exposure time was 1/8 second. Illumination consisted of two white light sources at both sides of the camera. In total, we took 508 images of 3872⍰×⍰2592 pixels with all external and internal side of fruits, and the label for each genotype in the same image.

### Preprocessing and segmentation

The first step in the pipeline is to segment and recognize the objects, since each raw image contains internal and external fruits, a rule, a coin and a printed genotype (the strawberry line) label. Image segmentation is needed for obtaining meaningful morphometric and color information. However, most of available technologies to determine the boundaries of the objects at the pixel level are usually semi-automatic and time-consuming [31–34]. Our fully automatic python-based pipeline takes the images of each strawberry line and outputs a curated database of square images (1000 px) and reads the genotype label (Fig. 1).

**Figure 1:**
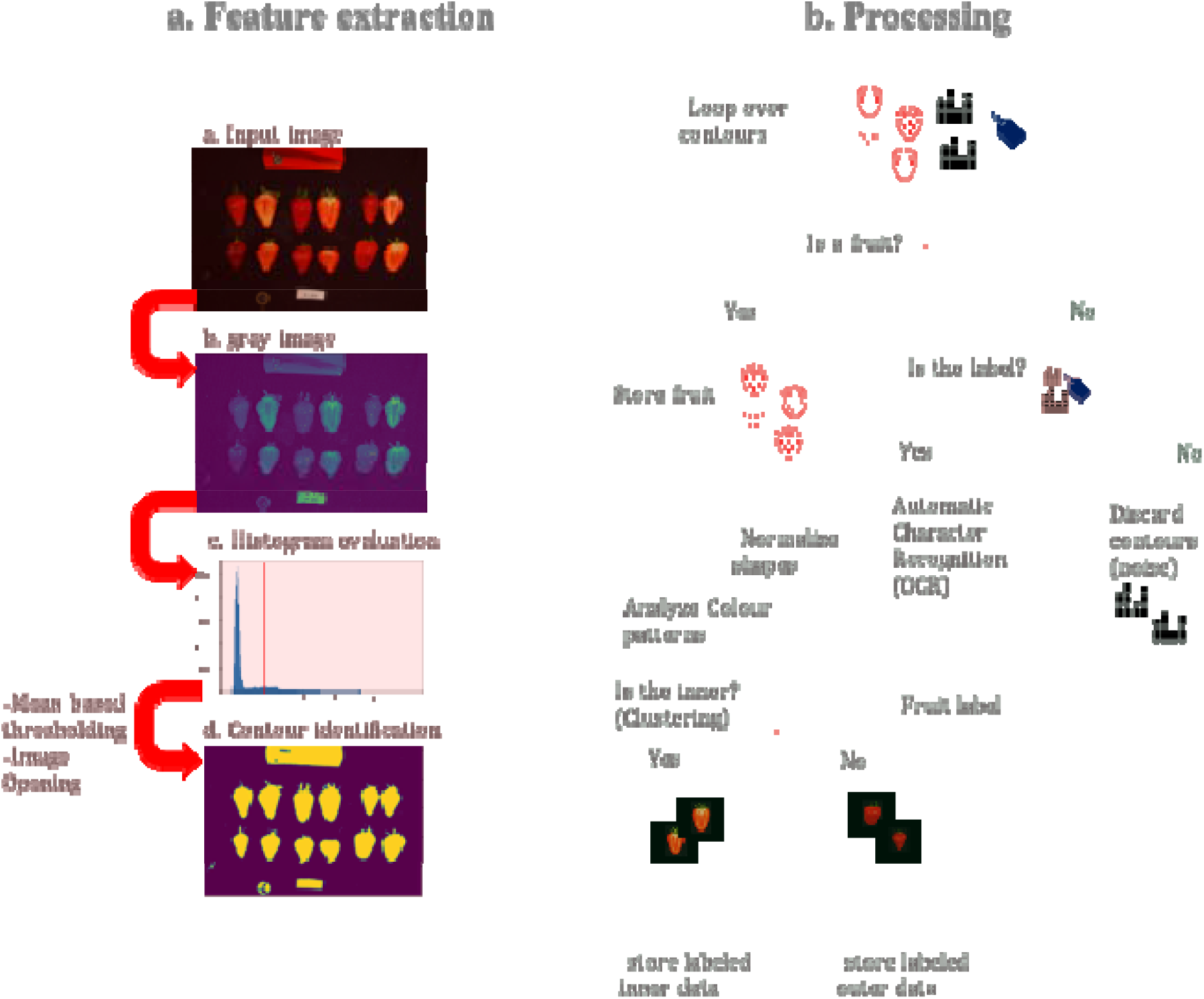
Workflow for automatic segmentation and label recognition from strawberry images. (a) Feature extraction part and (b) Feature preprocessing and database generation.

For segmentation, the three-channel digital signals (RGB/BGR) are converted into grayscale and blurred using Gaussian filtering of size 5, to remove undesirable noise. The histogram information is used for image binarization, i.e., splitting the background and foreground. Here, we binarized the image using just the mean value of the pixel as a threshold. The pipeline also allows the OTSU thresholding [35], which is designed to automatically define the threshold by minimizing the “overlap” between two classes. After binarization, we performed erosion and dilation, the former shrinks the edges and the later makes the image region grow. Finally, the algorithm extracts the Regions of Interests (ROI) and determines whether it is a strawberry or an image label. The color pattern analysis allows us to classify the internal or external part if a fruit image. For the labels, the Optical Character Recognition (OCR) algorithm from PyTesseract library (*https://pypi.org/project/pytesseract/*) is used to read the genotype name and automatically label the image into the database. As a result, the algorithm delivers a curated database of 508 folders labeled with the name of each genotype, and subfolders containing either the internal or external strawberry pictures (Fig. 1, Algorithm 1 in Suppl. Info).

### Automatic fruit phenotyping

Once masks for either internal or external fruit images are obtained, an automatic phenotyping procedure is run for inside or outside parts separately (Fig. 2). Classical linear descriptors, multivariate and deep learning techniques are combined from a novel perspective to dissect a variety shape and color patterns. If pedigree or marker information is available, a genetic analysis can be employed to estimate variance components for each of the fruit phenotypes. In the following, we describe the main methods implemented in the pipeline of Fig. 2.

**Figure 2.**
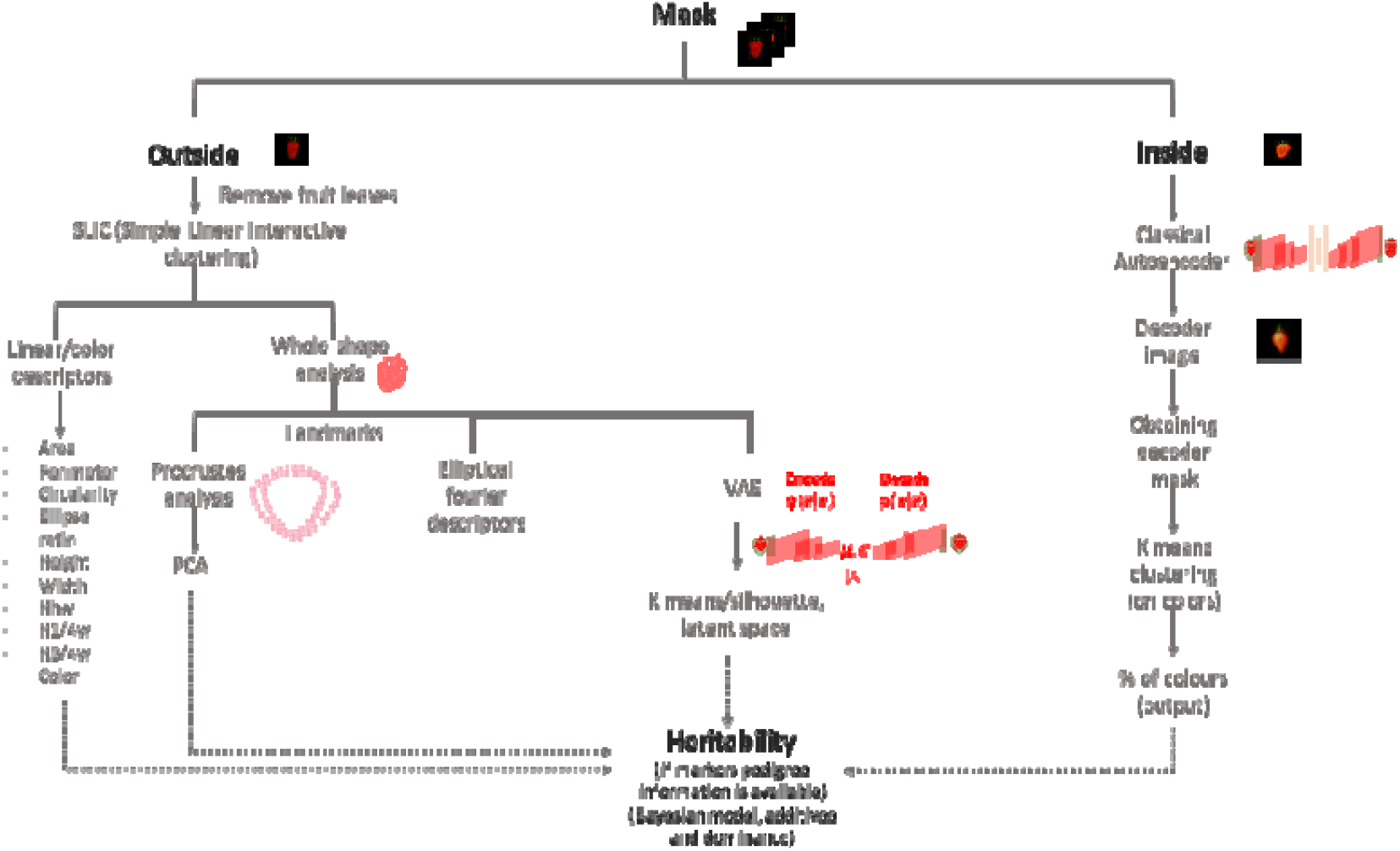
Data analysis workflow (available at https://github.com/lauzingaretti/DeepAFS). The input are all the segmented internal and external fruit images. External images are used for linear and multi-dimensional shape analysis through different standard and machine learning approaches, including deep learning. Inside images are used to estimate the color pattern of the internal fruit. Additive and dominant genetic components of each of the extracted morphometric and color phenotypes are finally estimated using Bayesian Linear Modeling using either pedigree or DNA marker information.

### Using an autoencoder and k-means to infer internal color patterns

We used an ‘autoencoder’ (AE) network to perform an unsupervised clustering of the internal images. An autoencoder (See Fig 3a.) is an unsupervised machine learning technique that applies backpropagation to train a neural network where the outputs are the same values as the inputs [36]. The AE gives new insight into image analysis by learning the structure about the data, i.e., it is not designed to copy an exact replicate of the input but instead to learn the repeatable and most useful properties.

**Figure 3.**
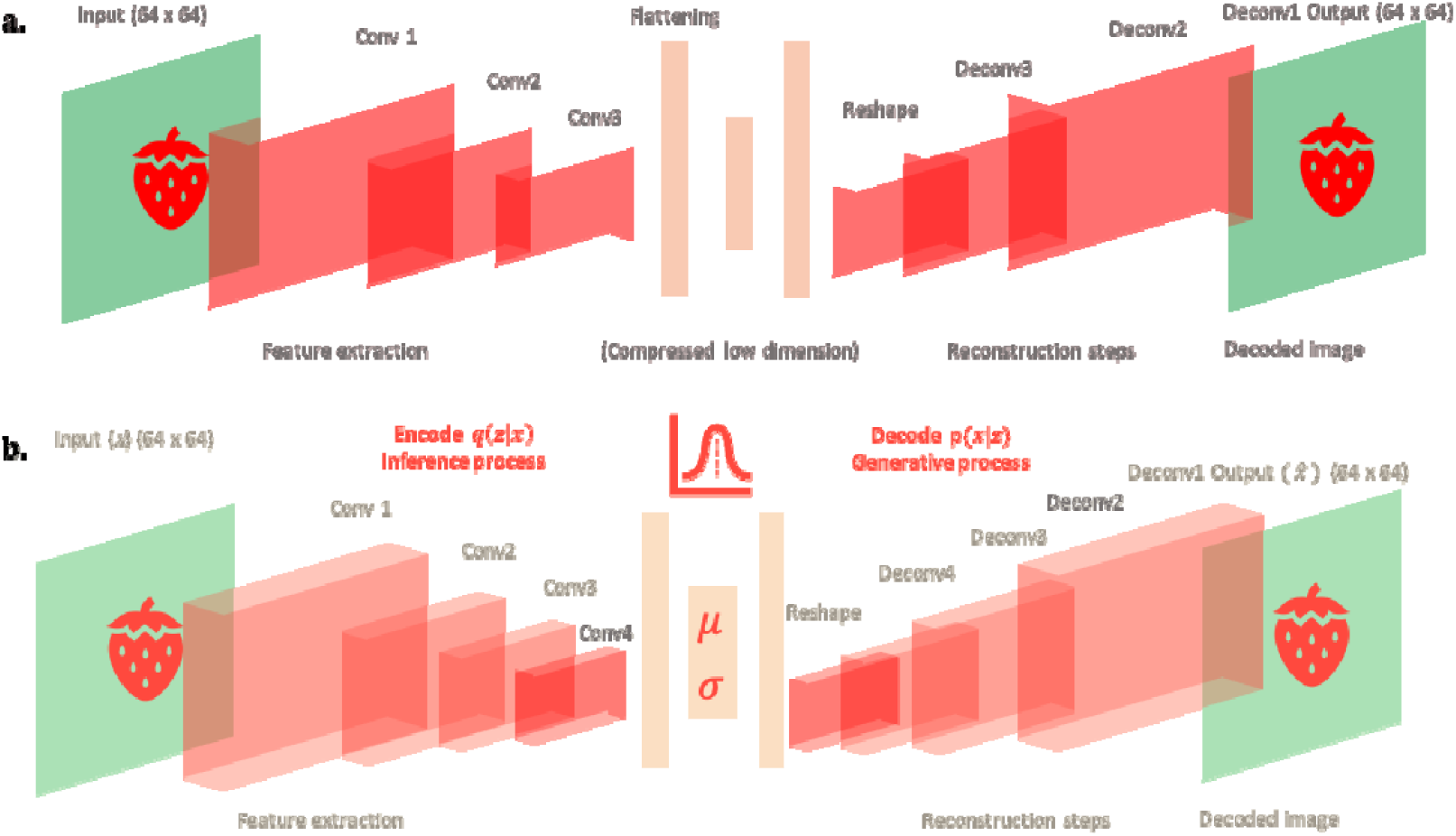
(a.) Architecture of convolutional autoencoder applied to the internal fruit images. (b) Architecture of convolutional variational autoencoder applied to external fruit. Unlike classical autoencoder, variational autoencoder are generative process as they learn the parameters of a distribution, instead of the feature representation. This network was trained using image of size 64×64, the encoder step consisted on 4 convolutional layers with kernel size equal to 3 and the linear rectified ‘relu’ as activation function to perform feature extraction, see details in github account. Finally, the convolution output is flattened, and the mean and sigma parameters are extracted from a dense layer. In the last network, the encoder step starts with a vector sampled from the latent distribution as input and reconstructs the input by performing deconvolution operations. The last deconvolution uses the sigmoid as activation function. The loss function is the Kulback-Leibler (KL) divergence, which consists of both, a ‘reconstruction’ and a ‘regularization’ term. The first network is a classical autoencoder, which uses the classical Mean Squared Error (MSE) as loss function.

We used a convolutional AE, as convolutional operations are especially suited for image analysis [36,37]. These layers create a feature map from the input image, preserving the relationships between pixels in the original space (Fig. 3a). Each convolution outputs a scored-filtered image, where a high score means a perfect match between the original and filtered image. The output layer is obtained by applying the Rectified Linear Unit (ReLU) activation function. Finally, as usual in any convolutional architecture, a max-pooling layer shrinks the output size and achieves a smoother representation, summarizing adjacent neuron outputs by computing their maximum.

The decoded images from an *AE* architecture are less noisy than the original ones, making it easier to detect repeatable/consistent color patterns. Our approach consists in taking 5 colors as reference: a class for the background (black) and four classes for the internal fruit color pattern, including leaves. The four ‘target classes’ “orange-like” (198, 99, 35, in RGB coordinates), “quasi-red” (184, 46, 8), “pale” (194, 144, 78), and “green” (76, 75, 20) for leaves. We then perform a k-means clustering with four classes after removing the background and we assigned each cluster to the nearest category among the four previously named using the Euclidean distance between the average color of each cluster and the references. As a result of this step, the surface of each of 1900 strawberry images is split in four categories of colors.

### Superpixel algorithm to remove leaves

Some of the fruit pictures had leaves. Leaves interfere with fruit shape quantification and need to be removed prior to estimating shape parameters. For that purpose, we applied the Simple Linear Interactive Clustering (SLIC) algorithm [38] from the Python skimage library. SLIC is based on the ‘superpixel’ concept. Basically, a superpixel is a group of pixels sharing perceptual and semantic information, e.g., the pixels in a superpixel are grouped together because of their color or texture features. The iterative algorithm starts with regularly spaced K-centers at a given distance, user defined as S, which are then relocated in the direction of the lowest gradient in a 3×3 neighborhood window to avoid being at the edges of the image. Further, a pixel is assigned to a given cluster if its distance to the cluster’s center is smaller than the distance to the other centers in the search area, as determined by S. Finally, the centers are recalculated by averaging all the pixels belonging to the superpixel. The iterative process ends when the residual error (distance between previous centers and recomputed ones) does not exceed a fixed threshold. SLIC outputs a set of meaningful clusters, splitting the background, the leaves and the fruit. Knowing that all our fruits are centered in the image, the superpixel containing the central pixel matches with the fruit.

### Univariate phenotypes: linear descriptors

Numerous object shape descriptors exist in the literature. Particularly for fruits, a controlled vocabulary was established in [23]. Here, we implement a custom script to compute some standard linear measures: circularity, solidity, shape aspect [25], ellipse ratio [23], fruit perimeter and area, fruit width at 25% height, fruit width at 75% height, fruit width at 50% of height, total height and maximum width. Circularity is a measure of the degree of roundness of a given object, defined as the ratio between the area of a given object and the area of a circle with the same convex perimeter, i.e., a value near one means a “globe” o “circular” shape. Solidity is the ratio between the area of the object and the area of the convex hull of a given shape. Most of the linear descriptors we used are standard in fruit shape analyses [23,25,33,39,40].

The external fruits color was estimated using the CIELAB space, were *L* indicates the luminosity, and *a* and *b* are the chromatic coordinates. The variation on the index *a* indicates the transition between green to red, where a higher value means a redder object. Variations in *b* reflect the change between yellow and blue color, i.e., a higher *b* value refers to a ‘bluer’ object

### Generalized Procrustes Analysis (GPA)

Shape is usually defined as all the geometric information that remains unchanged after filtering out the location, scale and rotation effects of a given object [27]. The above shape linear descriptors are standard in the literature but do not provide a whole shape portrayal. Alternatively to linear descriptors, shape variations can be described using ‘pseudo-landmarks’ [29], which identify points around the outline of the object. Fifty pseudo-landmarks were defined as the intersection between 50 equally spaced conceptual lines starting from the centroid and the fruit contour (Fig. 4a). Next we performed a Procrustes analysis [41]. The Procrustes analysis aims at finding the transformation T such that given two matrices *X*_*1*_, *X*_*2*_, the product *X*_*2*_ T best matches *X*_*1*_. The Generalized Procrustes Analysis (GPA) is an extension of the method devised to align many matrices simultaneously [41]. In morphometric analysis, this is done by averaging the distance between all the landmarks on a target shape and the corresponding points on a reference one. The pseudo-landmarks of the samples can then be analyzed as a multivariate object using, for instance, a principal component analysis (PCA). In addition, the pseudo-landmark variability gives insight on the most important regions that determine the differences between shapes. We used the Momocs [42] and geomorph [43] R packages to run these analyses.

**Figure 4.**
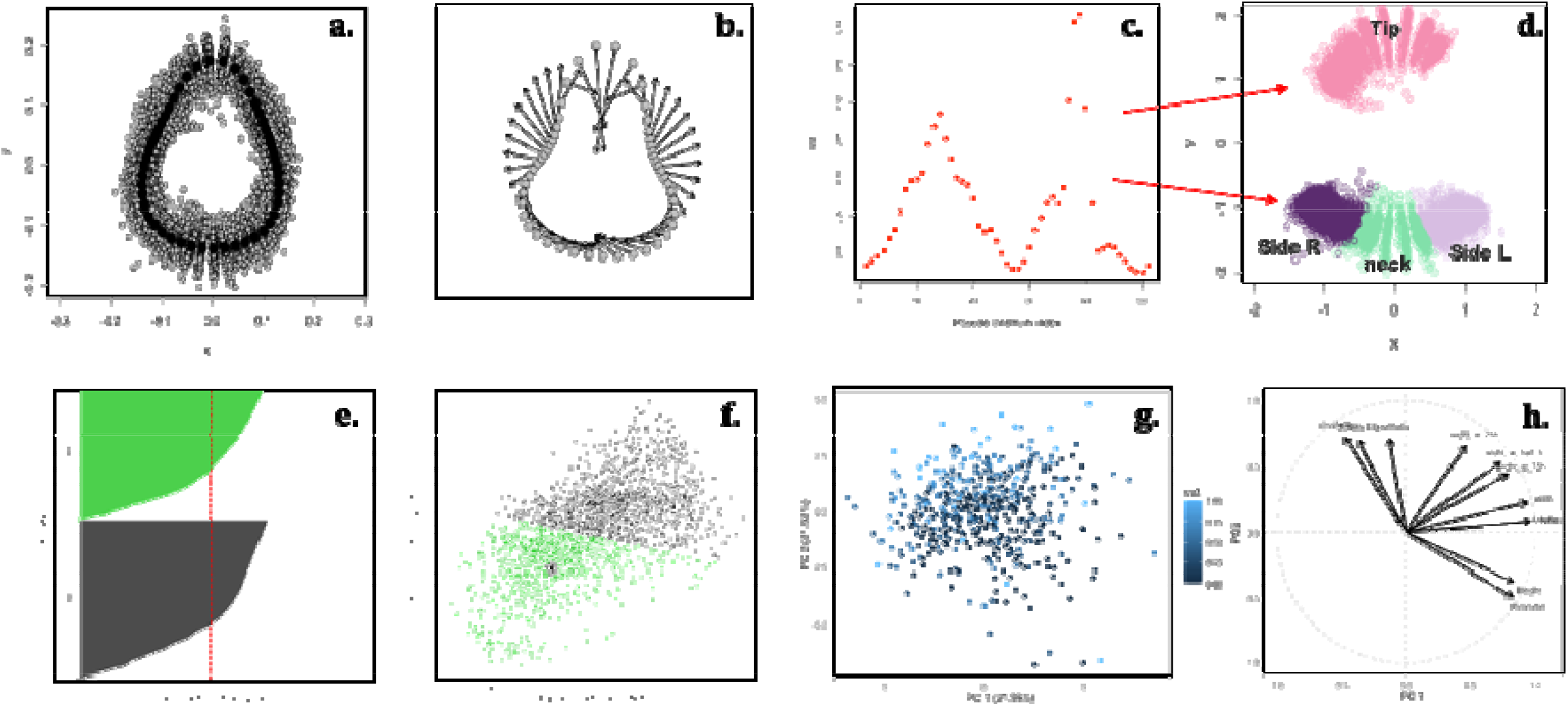
**(a)** Procrustes Analysis output: landmarking superimposition for all external fruit shapes. (b) Two extreme plot Procrustes analysis: mum and maximum consensus for the fruit shape. (c) standard deviation of each of the 50 landmarks, the average standard regression is the d line parallel to x-axis. Positions with a standard regression above the average are the most variable regions. (d) The most variable regions, h determines fruit shape are the tip, the neck and both sides around the neck. (e) Silhouette plot for k=2 clusters over the latent space (VAE), ilhouette for k=2 was 0.38. (f) Output for sample generation from VAE after applying k means with k=2. (g) PCA of all linear shape descriptors, dot represents a a different sampled and the color is proportional to the predicted proportion of fruit to each category from the clusters ned in (f). (h) Relationship between the linear shape variables from the PCA analysis.

### Elliptical Fourier Descriptors

An alternative approach to morphometric analysis is Elliptical Fourier transformation [44]. This method describes a closed curve as a sum of sine and cosine functions of growing frequencies. As its name suggests, Fourier harmonics are ellipses, and a larger number of harmonic means that more ellipses are fitted to a given contour. The second-order harmonic is simply one ellipse with the values of sine and cosine components for the x and y-axis, respectively. As the strawberry fruit is a relatively simple shape, four harmonics were enough to describe all the shapes in the database, giving a total of 16 coefficients. A PCA of the Fourier components can also be employed to quantify morphometric variability, as in procrustes analysis. Geomorph [43] R package was employed.

### Conditional Variational Autoencoders (VAE) to cluster shapes

Fruit shape can also be addressed from a completely different angle, such as obtaining clusters of shapes to objectively classify fruits in groups of similar morphology [39]. A standard approach consists of flattening the image and group the raw data, treating each pixel as a feature. Unfortunately, clustering algorithms are not exempt from the “curse of the dimensionality” problem [45] and they perform poorly as the number of analyzed dimensions increases, especially if noise is high.

A natural way to solve the aforementioned issue is to apply a dimension reduction technique before clustering. Although the classical autoencoders seem to be a good option, as shown above, AEs were conceived to perform a non-linear and not isometric dimensionality reduction, and thus they not preserve the geometrical properties of the original space [46]. Unlike traditional autoencoders, variational autoencoders [47–49] (VAEs) preserve distances and, importantly, are generative models (Fig. 3b). The main difference between *AE* and *VAE* is that the latter encodes the input as a distribution over a latent space. Basically, given an input x, VAE creates a latent distribution *p*(*z*|*x*) and the input reconstruction *d*(*z*) is obtained after sampling z from the latent representation *z*∼*p*(*z*|*x*). The VAE does not only force the latent space to be continuous, it can also generate meaningful information, even with images that it has never been seen before.

The key aspect in VAE training lies in the loss function, which includes a “reconstruction” and a “regularization” term. The former is the usual loss or the joint log-likelihood between the true and the VAE output, whereas the second is the entropy corresponding to the Kulback-Leibler divergence [36] between the latent distribution *N*(*μ*_*X*_, *σ*_*X*_) and the standard normal distribution *N*(*0,1*). Without incorporating a regularization, the VAE behaves as AE, where the latent space is neither complete nor continuous. Regularization forces the latent distribution to be close to the normal standard, generating a continuous space of low variance centered in the origin, which is suitable for data clustering and generation [36].

Here, we run standard k-means clustering of the latent space, with k varying between 2 and 9 groups. We chose a maximum k=9 given that up to nine strawberry shapes have been proposed in the literature, in particular in the Japanese market [50]. We assessed the cluster robustness using the silhouette index [51]. This index determines how well each object fits into its cluster, taking into account intra and between classes variations. The index ranges between −1 and 1, and a value close 1 means that the cluster is compact and homogeneous. Importantly, the combination of VAE and clustering also allows us to use conditional VAE to generate the expected fruit pertaining to a specific group.

### Genetic parameter inference

Genetic parameters determine how successful will be artificial selection and are therefore a critical parameter of any plant breeding scheme. Heritability (*h*^*2*^) is the proportion of phenotypic variance explained by the genetic variation [52]. To estimate it, we exploit the degree of resemblance between relatives using either pedigree or molecular information. Take linear modeL

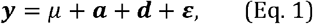

where ***y*** represents the phenotypes vector, averaged for each genotype, *μ* is the intercept, 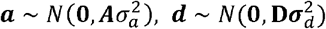, and 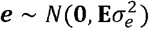 are the additive, dominant effects, and 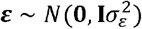 is the residual component, respectively; **A** = {a_ij_} and **D** = {d_ij_} are the additive and dominant covariance matrices, respectively. Both **A** and **D** can be computed recursively from the pedigree [53]. In the case of marker information, **A** and **D** can be computed as specified in [54,55], and implemented in [56] but statistical inference is otherwise identical. Posterior distributions of the genetic parameters were obtained using Reproducing Kernel Hilbert Spaces (RKHS) regression with the BGLR package [57]. The additive and dominance variance fraction were estimated as 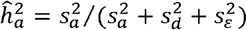 and 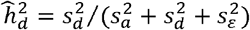, where 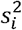 is the mean posterior estimate of 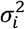.

## Results

### Shape descriptors

Shape linear descriptors, color in CIELAB scale, pseudo-landmarks and Elliptical Fourier transforms for fruit shape were computed for the 1920 external images output from Fig. 1 and Algorithm S1. Fig. 4b shows the minimum and maximum consensus for shape superimposition, suggesting that shapes vary between a “globose-like” to an “elongated-like” form in these samples. The standard deviation of the first PCA from GPA coordinates (Supp. Fig. 1) of tip, neck and both sides around the neck are above the mean (Fig. 4c). This suggests that these regions are responsible for the main shape variations in strawberry, in agreement with [39]. The Supp figure 1 shows the fruit shape variations from the Procrustes Principal Component Analysis (Proc-PCA). The first principal component describes the variations between ‘elongated’ to ‘globose’ like. Observations with a negative score on that components correspond to elongated fruits, while those who have positive scores are ‘globose’-like fruits. We then run a permutation-based Procrustes analysis of variance to assess the effect of the crosses on the fruit shape. The p-value obtained after 101 permutations shows a significant effect of the lines in the fruit shape (p<0.01), suggesting that the shape is heritable (Supp. Table 1).

We also set a fourth order elliptical Fourier to describe the main strawberry shape variations (see Supp. Figs. 2 and 3). As in the Procrustes Analysis, variations in the first principal component of the elliptical analysis show that the strawberry shapes vary between “globose-like” to “elongated-like”. Similarly, the first component from Elliptical -PCA analysis can also be used as a “morphological” descriptor. A k-means clustering using the two first PCA components of Fourier transform similarly detects the two previously defined groups of shapes setting k=2 (Supp. Fig. 4).

Alternatively, one can directly identify the number of different shapes from an image database. We used a VAE (Fig. 3b) to discover the optimal number of shapes in our database, which again was k=2 (Fig. 4 e,f, Supp. Fig. 5, 6). About 35% of the strawberries in our database belong to the ‘globose-like’ shape, whereas the remaining fruits were classified as ‘elongated-like’ (see Fig. 5a,b). The advantage of this analysis is that it can be used to automatize shape discovery in a completely unsupervised manner, and that is also capable of generating shapes not seen before, making it a powerful tool to predict shapes of new crosses or genotypes. As shape varies within and between genotypes, there are fruits of both classes for most of the genotypes; thus, we define a class shape index for each sample by calculating the proportion of fruits belonging to each category.

**Figure 5.**
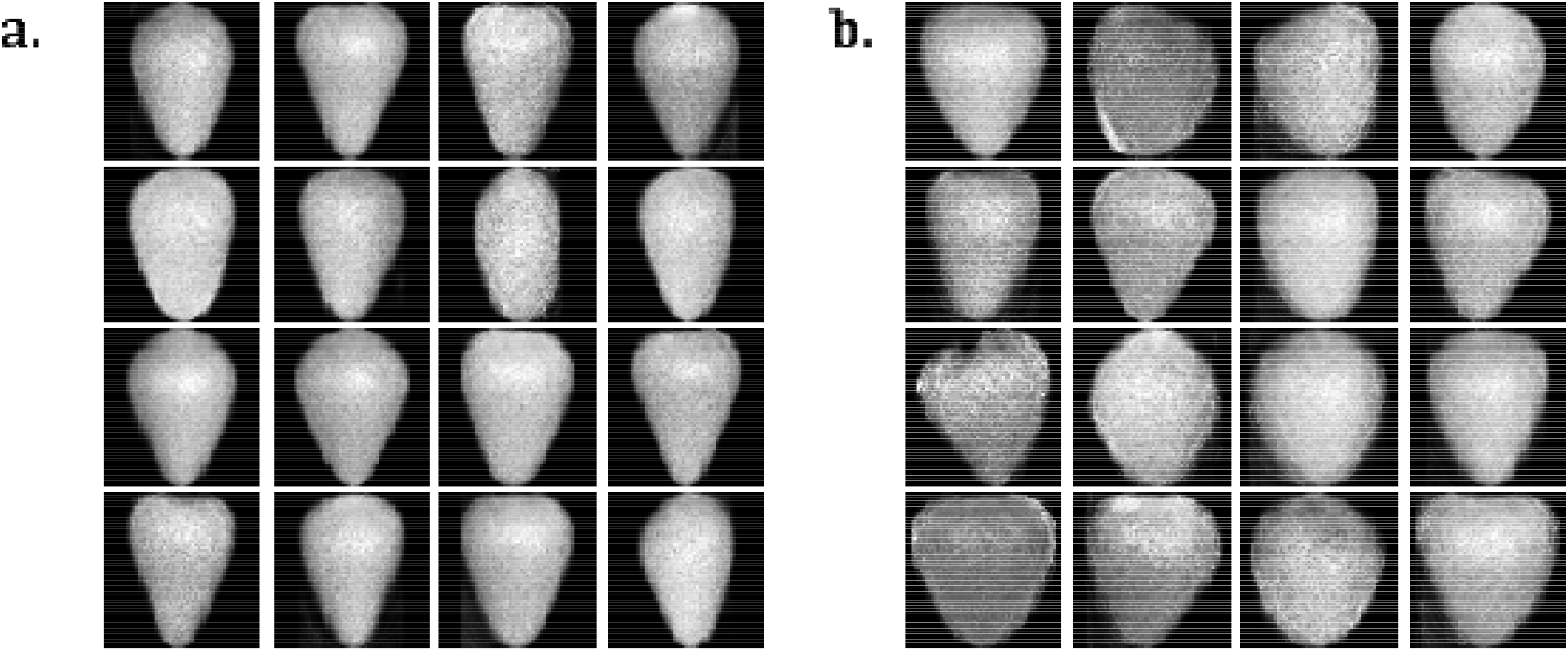
Images generated using the variational autoencoder combined with k-means in the latent space with k=2. (a) Images from the gated-like” cluster (b) Images from “globose-like” cluster.

**Figure 6.**
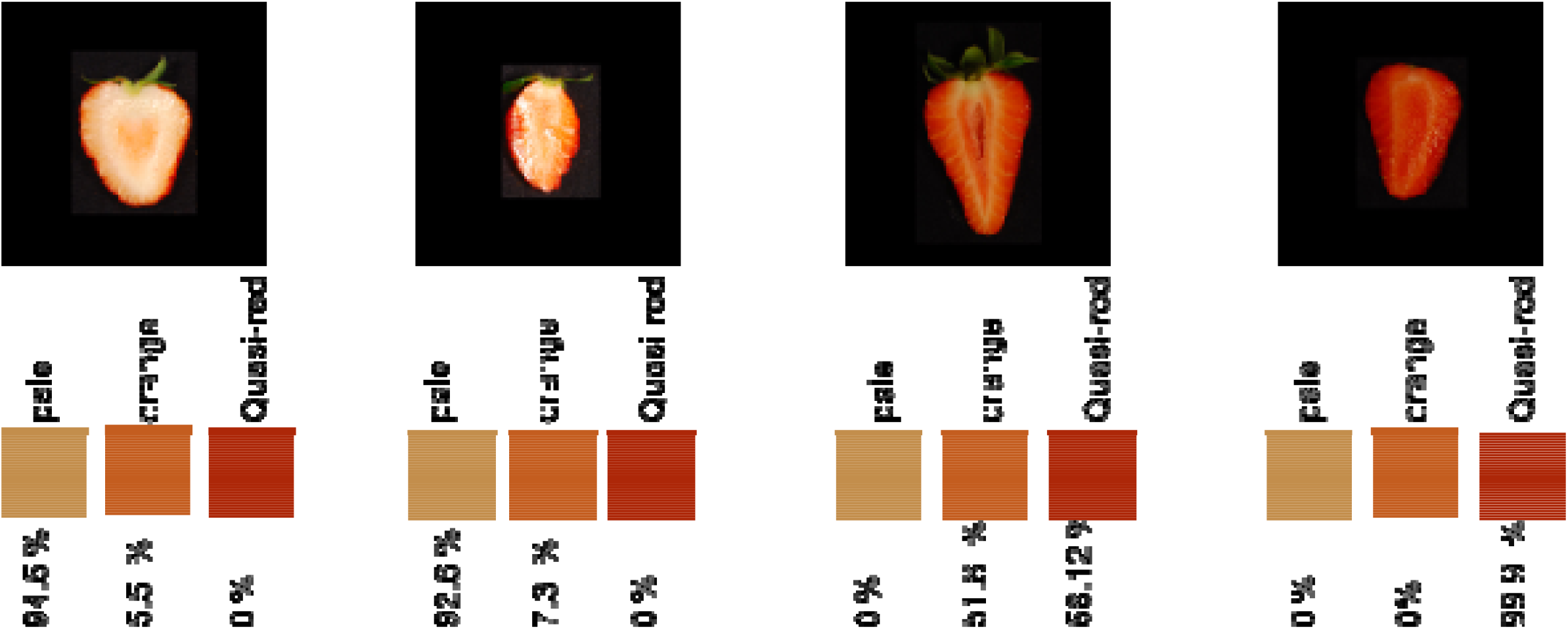
Estimated percentage of each of the three colors estimated for four chosen strawberries.

Fig. 4g shows a PCA on the linear descriptors, where the color of each sample is proportional to the predicted cluster probability. A dark color represents a fully elongated shape and light blue, a fully round fruit. Note that shape gradient is mainly observed along the second principal component. Interestingly, the most influential variables in this component are the fruit ratio between main and minor ellipse axis, the circularity and solidity coefficients (Fig. 4h). All of these are shape related variables. It is not surprising that solidity and circularity are highly correlated, since the convex hull area increases when a shape digresses from a circle (circularity), and solidity approaches zero. The area, perimeter and height are quasi-independent of the aforementioned descriptors and are not related with the shape clusters.

### Color descriptors

For the external side color in our dataset, *L* channel ranged between 7.01 and 118.30, mean of 75.54, *b* channel ranged between 127.9 and 184.8, mean value of 167.1, and *a* channel had a mean of 175.4, ranging between 128.8 and 192.6.

Estimating the color of the internal fruit is more challenging than that of external parts, as it fluctuates in a wider range of patterns. Fig. 6 shows the estimated percentages of each reference color for four chosen strawberries. Note the percentage of “quasi-red” is zero and most of the fruit is computed as “pale” (∼95%) for the first two, whitish fruits. Two colors, “quasi-red” and “orange-like”, predominate in the third fruit. Finally, the last fruit is almost red, as can be verified from the estimated quasi-red value (99%).

### Heritability estimation

Figure 7 shows the Bayesian estimates of heritability for all automatically extracted traits. We used the pedigree information to compute the additive and dominance relationship matrices, since we did not have SNP genotypes. Like many polyploid species, strawberry is clonally propagated [58]. Inferring the dominance component in these cases is critical, as clonal propagation allows a straightforward utilization of gene interaction [59]. Interestingly, we found that dominance variance was higher than the additive component for most of the traits. Broad sense heritability, i.e., the sum of both components ranged between 0.4 to 0.6, indicating than the traits are clearly heritable. The ellipse ratio, and the ratio between height and width were the most heritable characters, exhibiting an important additive component. Elliptical Fourier components, as well the percentage of fruits of each of both categories obtained from VAE also have a high heritability, for both additive and dominant components. Regarding the internal color, we find that the pale color has an important dominant component.

**Figure 7:**
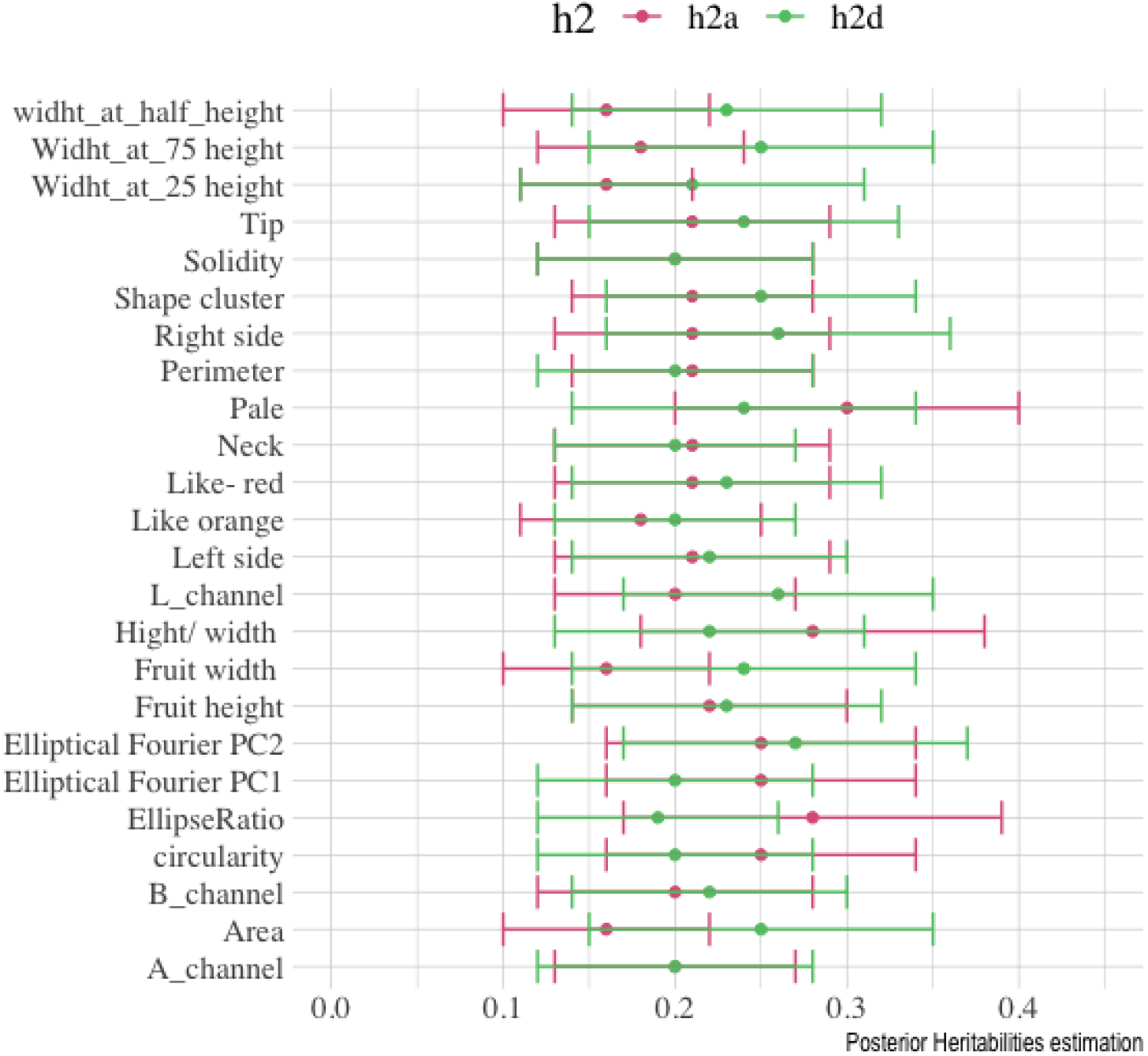
Estimation of additive (h2a) and dominance (h2d) variances for strawberry phenotypes.

## General Discussion

Over the last decades, plant and animal breeding programs have benefited from the development and cost reduction on genomic technologies [60,61]. Breeding nevertheless depends of both genotype and phenotype, and our ability of characterizing the latter is much more limited compared to the former [10,62]. In fact, one of the biggest challenges of ‘Precision Agriculture’ is to transform large-scale datasets collected with sensors into phenotypic measurements that can be used for genetic improvement.

Consumer attitudes are increasingly shaping agricultural practices. In the case of fruits, consumer preferences are based primarily on fruit appearance. However, measuring this trait is not straightforward, as it is a complex mixture of shape and color patterns. A crucial aspect for improving appearance is then to characterize the color and shape of the fruits in an inexpensive and fast way. In this paper, we deliver a fully automatized pipeline that analyzes fruit appearance as complex multivariate data. While this is not the first study characterizing fruit shape variations, our procedure is quite more automatized than their predecessors as requires minimal human intervention [23,34,39]. It also incorporates new features such as the use of variational autoencoders (VAE) to automatically detect the most likely number of underlying shapes or to clustering the internal color.

Digital images are among the easiest phenotypes to collect. Analyzing them is nevertheless challenging, partly because objects boundaries must be determined, a process known as feature extraction. Numerous classical [35,38,63] and deep learning approaches [64,65] have been developed in computer vision and image processing to meet this objective. Here, we combined some of these methods to automatically segment fruit snapshots and read the fruit label. The main approach we used is not new, as it is based in an algorithm developed in the late seventies [35]. However, we resort to novel techniques in order to remove undesirable image noise [66], and we characterize color pattern or classify fruits through a variational autoencoder [36].

The phenotype results from a complex interaction between the genotype and environmental factors. In this work, we characterize shape and color variations using several complementary methods, from naÏve linear descriptors to multivariate and deep learning techniques. It is important to point out that results from all approaches are consistent, and suggest that the fruits in our database can be classified in two groups, “globose-like” and “elongated-like” (Figs. 4e,f, 5). We determine that the most variable regions are the neck, neck-sides and the tip of the fruit (Fig. 4c,d). The “shape” linear descriptor, i.e., the ratio between fruit shape and height, is a good morphological descriptor (Fig. 4g,h) and is as discriminative as more complex multivariate characterizations. An ANOVA on the Procrustes coordinates shows that genotype is significant (p-value<0.01, Supp Table 1), another indirect indication that shape is inheritable.

Describing internal color patterns is challenging, mainly because color is a quantitative multi-channel character. We addressed this problem by defining three reference colors named as “quasi-red”, “orange-like” and “pale”. We then automatically determined the percentage of color corresponding to each of these target classes for each fruit using an autoencoder for fruit denoising and a k-means for segmentation. The algorithm calculates the Euclidean distance between the three RGB coordinates obtained by means of clustering to the target color coordinates and classifies the cluster as belonging to one of the three targets whose distance is minimal. The colors patterns are satisfactorily dissected, as can be seen in some picked images from the database (Fig. 6.).

Portraying the phenotypes would not be worthwhile for breeding if the desirable characters could not be transmitted to the progeny. Thus, quantifying the heritability of all of these traits is crucial. Typically, genetic variance is decomposed in additive and non-additive effects [67]. Clonally propagated species like strawberry allows direct utilization of dominance and epistatic interaction. We used Bayesian modeling to estimate both additive and dominant effects. As can be observed in Fig. 7 and Supp. Table 2, most traits are moderately heritable, and a high degree of variance is explained by the dominance component. In this scenario, prediction accuracy can be significantly increased in genomic selection by including dominance in the model [58].

We estimated heritabilities using pedigree information, but a similar study can be carried out if genetic markers are available. This would also have an extra benefit, as markers allows to perform Genome Wide Association studies (GWAS) and to implement genomic selection. It is straightforward to implement these features in our pipeline. Association studies for human shape variations, apple leaf or quantitative analysis for tomato have revealed genes or markers associated with craniofacial shape [28,68,69], leaf variation [70] and tomato morphology [71]. To the best of our knowledge, there is not a similar study in strawberry and there is still a long way to go to unravel the genetic basis of strawberry shape [72].

Overall, our results show that, although fruit shape is made up of a complex set of traits, it can be quickly and automatically evaluated and is moderately heritable (Figs. 1,2,7). Although we focused in strawberries, the pipeline can be employed for high-throughput image phenotype of other fruit species, as shown by a few examples in GitHub (https://github.com/lauzingaretti/DeepAFS). Future improvements are still needed as, e.g., image segmentation is not always that simple. Future improvements should also address additional technological developments such as spectral and MIR images [17]. Finally, we underscore the need to develop analysis pipelines for plant high-throughput phenotyping, suitable to automate processes that are often subjective and time consuming.

## Acknowledgments

The authors would like to thank Planasa for providing the strawberry fruit under the Planasa-Irta collaboration contract. LMZ was supported by a PhD grant from the Ministry of Economy and Science (MINECO, Spain), work funded by the MINECO grants AGL2016-78709-R and PID2019-108829RB-I00 to MPE and from the EU through the BFU2016-77236-P (MINECO/AEI/FEDER, EU) and the “Centro de Excelencia Severo Ochoa 2016-2019” award SEV-2015-0533.

## Author Contributions

LMZ, AM and MPE conceived research. AM provided data. LMZ developed methods and code. LMZ and MPE wrote the manuscript with help from AM.

## Data Availability

Code is available at https://github.com/lauzingaretti/DeepAFS.

## Bibliography

1. Hunter MC, Smith RG, Schipanski ME, Atwood LW, Mortensen DA. Agriculture in 2050: Recalibrating targets for sustainable intensification [Internet]. Bioscience. Oxford University Press; 2017 [cited 2020 Sep 22]. p. 386–91. Available from: https://pennstate.pure.elsevier.com/en/publications/agriculture-in-2050-recalibrating-targets-for-sustainable-intensi

2. Carnicer J, Coll M, Ninyerola M, Pons X, Sánchez G, Peñuelas J. Widespread crown condition decline, food web disruption, and amplified tree mortality with increased climate change-type drought. Proc Natl Acad Sci U S A. 2011;108:1474–8.

3. Wernberg T, Smale DA, Tuya F, Thomsen MS, Langlois TJ, De Bettignies T, et al. An extreme climatic event alters marine ecosystem structure in a global biodiversity hotspot. Nat Clim Chang. Nature Publishing Group; 2013;3:78–82.

4. Hegerl GC, Hanlon H, Beierkuhnlein C. Climate science: Elusive extremes. Nat Geosci. Nature Publishing Group; 2011;4:142–3.

5. Porfirio LL, Newth D, Finnigan JJ, Cai Y. Economic shifts in agricultural production and trade due to climate change. Palgrave Commun [Internet]. Palgrave Macmillan Ltd.; 2018 [cited 2020 Sep 22];4:1–9. Available from: https://www.nature.com/articles/s41599-018-0164-y

6. Lobos GA, Camargo A V., del Pozo A, Araus JL, Ortiz R, Doonan JH. Editorial: Plant Phenotyping and Phenomics for Plant Breeding. Front Plant Sci [Internet]. Frontiers Media S.A.; 2017 [cited 2020 Sep 22];8:2181. Available from: http://journal.frontiersin.org/article/10.3389/fpls.2017.02181/full

7. Pieruschka R, Schurr U. Plant Phenotyping: Past, Present, and Future. Plant Phenomics [Internet]. 2019 [cited 2020 Sep 18];2019:1–6. Available from: https://spj.sciencemag.org/plantphenomics/2019/7507131/

8. Fasoula DA, Fasoula VA. Gene Action and Plant Breeding. Plant Breed Rev. John Wiley & Sons, Inc.; 2010. p. 315–74.

9. Tardieu F, Cabrera-Bosquet L, Pridmore T, Bennett M. Plant Phenomics, From Sensors to Knowledge. Curr. Biol. Cell Press; 2017. p. R770–83.

10. Awada L, Phillips PWB, Smyth SJ. The adoption of automated phenotyping by plant breeders. Euphytica. Springer Netherlands; 2018;214.

11. Mahlein A-K. Present and Future Trends in Plant Disease Detection. Plant Dis. 2016;100:1–11.

12. Stafford J V. Implementing precision agriculture in the 21st century. J Agric Eng Res. 2000;76:267–75.

13. Schork NJ. Genetics of Complex Disease Approaches, Problems, and Solutions. Am J Respir Crit Care Med. 1997.

14. Großkinsky DK, Svensgaard J, Christensen S, Roitsch T. Plant phenomics and the need for physiological phenotyping across scales to narrow the genotype-to-phenotype knowledge gap [Internet]. J. Exp. Bot. Oxford University Press; 2015 [cited 2020 Sep 18]. p. 5429–40. Available from: https://academic.oup.com/jxb/article/66/18/5429/482901

15. Zhao C, Zhang Y, Du J, Guo X, Wen W, Gu S, et al. Crop phenomics: Current status and perspectives. Front. Plant Sci. Frontiers Media S.A.; 2019.

16. Gracia-Romero A, Kefauver SC, Fernandez-Gallego JA, Vergara-Díaz O, Nieto-Taladriz MT, Araus JL. UAV and Ground Image-Based Phenotyping: A Proof of Concept with Durum Wheat. Remote Sens [Internet]. MDPI AG; 2019 [cited 2020 Sep 22];11:1244. Available from: https://www.mdpi.com/2072-4292/11/10/1244

17. Ruckelshausen A, Busemeyer L. Toward digital and image-based phenotyping. Phenomics Crop Plants Trends, Options Limitations. 2015.

18. Jaeger SR, Machín L, Aschemann-Witzel J, Antúnez L, Harker FR, Ares G. Buy, eat or discard? A case study with apples to explore fruit quality perception and food waste. Food Qual Prefer. Elsevier Ltd; 2018;69:10–20.

19. Gilbert JL, Olmstead JW, Colquhoun TA, Levin LA, Clark DG, Moskowitz HR. Consumer-assisted selection of blueberry fruit quality traits. HortScience [Internet]. American Society for Horticultural Science; 2014 [cited 2020 Sep 23];49:864–73. Available from: http://www.

20. Lewers KS, Newell MJ, Park E, Luo Y. Consumer preference and physiochemical analyses of fresh strawberries from ten cultivars. Int J Fruit Sci [Internet]. Taylor and Francis Inc.; 2020 [cited 2020 Sep 23];1–24. Available from: https://www.tandfonline.com/doi/full/10.1080/15538362.2020.1768617

21. González M, Baeza E, Lao JL, Cuevas J. Pollen load affects fruit set, size, and shape in cherimoya. Sci Hortic (Amsterdam). Elsevier; 2006;110:51–6.

22. Klatt BK, Holzschuh A, Westphal C, Clough Y, Smit I, Pawelzik E, et al. Bee pollination improves crop quality, shelf life and commercial value. Proc R Soc B Biol Sci [Internet]. Royal Society; 2014 [cited 2020 Sep 23];281:20132440. Available from: https://royalsocietypublishing.org/doi/10.1098/rspb.2013.2440

23. Brewer MT, Lang L, Fujimura K, Dujmovic N, Gray S, Van Der Knaap E. Development of a controlled vocabulary and software application to analyze fruit shape variation in tomato and other plant species. Plant Physiol [Internet]. American Society of Plant Biologists; 2006 [cited 2020 Oct 5];141:15–25. Available from: www.plantphysiol.org/cgi/doi/10.1104/pp.106.077867.

24. Brewer MT, Moyseenko JB, Monforte AJ, Van Der Knaap E. Morphological variation in tomato: A comprehensive study of quantitative trait loci controlling fruit shape and development. J Exp Bot. 2007;58:1339–49.

25. Rashidi M, Keshavarzpour F. Classification of Apple Size and Shape Based on Mass and Outer Dimensions. J Agric Environ Sci [Internet]. 2010 [cited 2020 Oct 5];9:618–21. Available from: http://faostat.fao.org.

26. Mezghani N, Zaouali I, Amri WB, Rouz S, Simon PW, Hannachi C, et al. Fruit morphological descriptors as a tool for discrimination of Daucus L. germplasm. Genet Resour Crop Evol. 2014;61:499–510.

27. Peter Klingenberg C. Evolution and development of shape: integrating quantitative approaches. Nat Publ Gr [Internet]. 2010 [cited 2020 Jun 22]; Available from: www.nature.com/reviews/genetics

28. Claes P, Liberton DK, Daniels K, Rosana KM, Quillen EE, Pearson LN, et al. Modeling 3D Facial Shape from DNA. PLoS Genet. 2014;10.

29. Dryden, Ian L., Mardia K V. Statistical shape analysis. Wiley series in probability and statistics; 1998.

30. UPOV. Strawberry: Guidelines for the conduct of tests for distinctness, uniformity and stability. Upov. 2012;1–26.

31. Schindelin J, Rueden CT, Hiner MC, Eliceiri KW. The ImageJ ecosystem: An open platform for biomedical image analysis. Mol Reprod Dev [Internet]. John Wiley & Sons, Ltd; 2015 [cited 2020 Oct 1];82:518–29. Available from: https://onlinelibrary.wiley.com/doi/full/10.1002/mrd.22489

32. Schindelin J, Arganda-Carreras I, Frise E, Kaynig V, Longair M, Pietzsch T, et al. Fiji: An open-source platform for biological-image analysis. Nat. Methods. 2012. p. 676–82.

33. Darrigues A, Hall J, Van Der Knaap E, Francis DM, Dujmovic N, Gray S. Tomato analyzer-color test: A new tool for efficient digital phenotyping. J Am Soc Hortic Sci. 2008;133:579–86.

34. Gehan MA, Fahlgren N, Abbasi A, Berry JC, Callen ST, Chavez L, et al. PlantCV v2: Image analysis software for high-throughput plant phenotyping. PeerJ [Internet]. PeerJ Inc.; 2017 [cited 2020 Oct 1];2017:e4088. Available from: http://plantcv.danforthcenter.org/pages/data.html

35. Otsu N. THRESHOLD SELECTION METHOD FROM GRAY-LEVEL HISTOGRAMS. IEEE Trans Syst Man Cybern. 1979;SMC-9:62–6.

36. Goodfellow, I., Bengio, Y., Courville A. Deep Learning. MIT Press. MIT Press Cambridge; 2016.

37. LeCun Y, Bengio Y, Hinton G. Deep learning. Nature. 2015;521:436–44.

38. Achanta R, Shaji A, Smith K, Lucchi A, Fua P, Süsstrunk S. SLIC Superpixels.

39. Feldmann MJ, Hardigan MA, Famula RA, López CM, Tabb A, Cole GS, et al. Multi-dimensional machine learning approaches for fruit shape phenotyping in strawberry. Gigascience. 2020;9:1–17.

40. Diaz-Garcia L, Covarrubias-Pazaran G, Schlautman B, Grygleski E, Zalapa J. Image-based phenotyping for identification of QTL determining fruit shape and size in American cranberry (Vaccinium macrocarpon L.). PeerJ. 2018;2018:1–19.

41. Gower JC. Generalized procrustes analysis. Psychometrika. 1975;40:33–51.

42. Bonhomme V, Picq S, Gaucherel C, Claude J. Momocs: Outline analysis using R. J Stat Softw. 2014;56:1–24.

43. Adams DC, Otárola-Castillo E. Geomorph: An r package for the collection and analysis of geometric morphometric shape data. Methods Ecol Evol [Internet]. John Wiley & Sons, Ltd; 2013 [cited 2020 Oct 15];4:393–9. Available from: https://besjournals.onlinelibrary.wiley.com/doi/full/10.1111/2041-210X.12035

44. Kuhl FP, Giardina CR. Elliptic fourier features of a closed contour. Comput Graph Image Process. 1982;18:236–58.

45. Bellman R. Dynamic Programming Princeton University Press [Internet]. Princeton, NJ. 1957 [cited 2020 Oct 13]. Available from: https://press.princeton.edu/books/paperback/9780691146683/dynamic-programming

46. Gropp A, Atzmon M, Lipman Y. ISOMETRIC AUTOENCODERS. arXiv Prepr 200609289. 2020;

47. Kingma DP, Welling M. Auto-encoding variational bayes. 2nd Int Conf Learn Represent ICLR 2014 - Conf Track Proc. 2014.

48. Kingma DP. Fast Gradient-Based Inference with Continuous Latent Variable Models in Auxiliary Form. 2013 [cited 2020 Oct 13]; Available from: http://arxiv.org/abs/1306.0733

49. Rezende DJ, Mohamed S, Wierstra D. Stochastic Back-propagation and Variational Inference in Deep Latent Gaussian Models. Proc 31st … [Internet]. 2014 [cited 2020 Oct 13];32:1278–86. Available from: http://jmlr.org/proceedings/papers/v32/rezende14.html

50. Ishikawa T, Hayashi A, Nagamatsu S, Kyutoku Y, Dan I, Wada T, et al. Classification of strawberry fruit shape by machine learning. Int Arch Photogramm Remote Sens Spat Inf Sci - ISPRS Arch [Internet]. 2018 [cited 2020 Oct 13]. p. 463–70. Available from: https://doi.org/10.5194/isprs-archives-XLII-2-463-2018

51. Rousseeuw PJ. Silhouettes: A graphical aid to the interpretation and validation of cluster analysis. J Comput Appl Math. 1987;20:53–65.

52. Falconer, Douglas S and Mackay Trudy FC and Frankham R. Introduction to quantitative genetics (4th edn). Trends Genet. 1996;12:280.

53. Nazarian A, Gezan SA. GenoMatrix: A Software Package for Pedigree-Based and Genomic Prediction Analyses on Complex Traits. J Hered [Internet]. Oxford University Press; 2016 [cited 2020 Oct 13];107:372–9. Available from: /pmc/articles/PMC4888442/?report=abstract

54. VanRaden PM. Efficient methods to compute genomic predictions. J Dairy Sci. Elsevier Inc.; 2008;91:4414–23.

55. Vitezica ZG, Varona L, Legarra A. On the Additive and Dominant Variance and Covariance of Individuals Within the Genomic Selection Scope. Genetics [Internet]. 2013 [cited 2019 Oct 11];195:1223–30. Available from: http://www.ncbi.nlm.nih.gov/pubmed/24121775

56. Amadeu RR, Cellon C, Olmstead JW, Garcia AAF, Resende MFR, Muñoz PR, et al. AGHmatrix: R Package to Construct Relationship Matrices for Autotetraploid and Diploid Species: A Blueberry Example. 2016; Available from: https://github.com/prmunoz/AGHmatrix/blob/master/

57. Pérez P, De Los Campos G. Genome-wide regression and prediction with the BGLR statistical package. Genetics [Internet]. 2014;198:483–95. Available from: http://www.ncbi.nlm.nih.gov/pubmed/25009151

58. Gezan SA, Osorio LF, Verma S, Whitaker VM. An experimental validation of genomic selection in octoploid strawberry. Hortic Res [Internet]. 2017;4:16070. Available from: http://www.nature.com/articles/hortres201670

59. Grüneberg W, Mwanga R, Andrade M, Espinoza J. Selection methods. Part 5: Breeding clonally propagated crops. Plant Breed Farmer Particip [Internet]. 2009;275–322. Available from: http://www.cabdirect.org/abstracts/20103075062.html

60. Robertsen CD, Hjortshøj RL, Janss LL. Genomic selection in cereal breeding. Agronomy. MDPI AG; 2019.

61. J. Crossa, P. Pérez-Rodríguez, J. Cuevas, O. Montesinos-López, D. Jarquín, Campos G de los. Genomic selection in plant breeding: Methods, models, and perspectives. Trends Plant Sci. 2017;22:961–75.

62. Tardieu F, Cabrera-Bosquet L, Pridmore T, Bennett M. Plant Phenomics, From Sensors to Knowledge. Curr. Biol. Cell Press; 2017. p. R770–83.

63. Unit MI. A review on image segmentation techniques. 1993;26.

64. He K, Gkioxari G, Dollár P, Girshick R. Mask R-CNN.

65. Badrinarayanan V, Kendall A, Cipolla R. SegNet: A Deep Convolutional Encoder-Decoder Architecture for Image Segmentation. IEEE Trans Pattern Anal Mach Intell [Internet]. 2017 [cited 2019 Oct 11];39:2481–95. Available from: https://ieeexplore.ieee.org/document/7803544/

66. Mao X-J, Shen C, Yang Y-B. Image Restoration Using Convolutional Auto-encoders with Symmetric Skip Connections. 2016;1–17. Available from: http://arxiv.org/abs/1606.08921

67. Zingaretti LM, Gezan SA, Ferrão LF V., Osorio LF, Monfort A, Muñoz PR, et al. Exploring Deep Learning for Complex Trait Genomic Prediction in Polyploid Outcrossing Species. Front Plant Sci [Internet]. 2020;11:1–14. Available from: https://www.frontiersin.org/article/10.3389/fpls.2020.00025/full

68. Claes P, Roosenboom J, White JD, Swigut T, Sero D, Li J, et al. Genome-wide mapping of global-to-local genetic effects on human facial shape. Nat Genet [Internet]. Springer US; 2018;50:414–23. Available from: http://dx.doi.org/10.1038/s41588-018-0057-4

69. Galvánek M, Furmanová K, Chalás I, Sochor J. Automated facial landmark detection, comparison and visualization. Proc - SCCG 2015 31st Spring Conf Comput Graph. 2015;7–14.

70. Migicovsky Z, Li M, Chitwood DH, Myles S. Morphometrics reveals complex and heritable apple leaf shapes. Front Plant Sci. 2018;8:1–14.

71. Rodríguez GR, Muños S, Anderson C, Sim SC, Michel A, Causse M, et al. Distribution of SUN, OVATE, LC, and FAS in the tomato germplasm and the relationship to fruit shape diversity. Plant Physiol. 2011;156:275–85.

72. Gaston A, Osorio S, Denoyes B, Rothan C. Applying the Solanaceae Strategies to Strawberry Crop Improvement. Trends Plant Sci. 2020;25:130–40.

