## Supplementary material for "Automatic fruit morphology phenome and genetic analysis: An application in the octoploid strawberry": Supplemenrary material

**SUPPLEMENTARY MATERIAL:**

**Algorithm Sup 1:**

This is the pseudo-code used for automatic morphology analysis for strawberry images. The code can analyze any fruit shape in similar conditions (homogeneous background) and it can be easily extended for custom purposes.

**Algorithm 1: Create a segmented fruit database from raw data**

n: number of images to process.

**for** i=1 to n **do**:

1. Read image
2. Convert RGB/BGR image into a grayscale image.
3. Smooth image using Gaussian filtering
4. Binarize image using mean based adaptative thresholding methodology
5. Apply erosion + dilation (Opening operation)
6. To obtain image contours

sh=[] (empty list)

**for** c in contours **do**:

1. Obtain *h, w* (contour height and width)
2. Obtain x, y (contour position in the main image)
3. **If** 1.1<(h/w)<3 (What is the expected aspect for fruits?) :

sh+=c

Analyze sh color pattern (to determine whether it is the inside or outside of the fruits) (Skip this step if your fruits are all inner (outer))

4. Get the ROI (Region of interest) from the RGB/BGR image.
5. Create an equal size image for each contour
7. Use OCR (Optical character recognition) to read the image label
8. Output a folder named by (7) containing the sample images (it splits inner/outer if necessary)

Algorithm available at <https://github.com/lauzingaretti>/DeepAFS

**Procrustes PCA:** Procrustes Principal Component analysis (Proc-PCA) on fruit shape. We evaluated the effect of the crosses on the fruit shape through a Procrustes analysis of variance using residual randomization permutation procedure with 101 permutations.

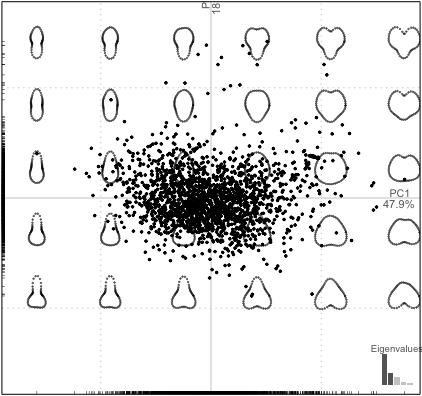

**Supp Fig. 1** Output plot from Procrustes Principal Component Analysis (Proc-PCA). The analysis shows a variation between ‘elongated’ and ‘globose’-like shape.

```
            Df     SS        MS     Rsq      F      Z.        Pr(>F)
crosses        24 0.5388 0.0224483 0.05525 4.5615 7.4238      0.009901 **
Residuals 1872 9.2126 0.0049213 0.94475
Total     1896 9.7514
---
Signif. codes:
0 ‘***’ 0.001 ‘**’ 0.01 ‘*’ 0.05 ‘.’ 0.1 ‘ ’ 1
```

**Supp Table 1.** Output from Procrustes ANOVA, which evaluates the effect of the crosses on fruit shape.

**Fourier Analysis**

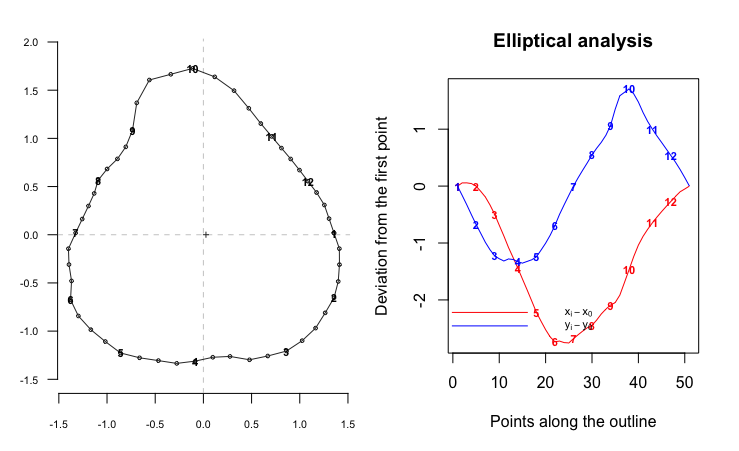

**Supp Fig. 2.** Output from elliptical Fourier analysis.

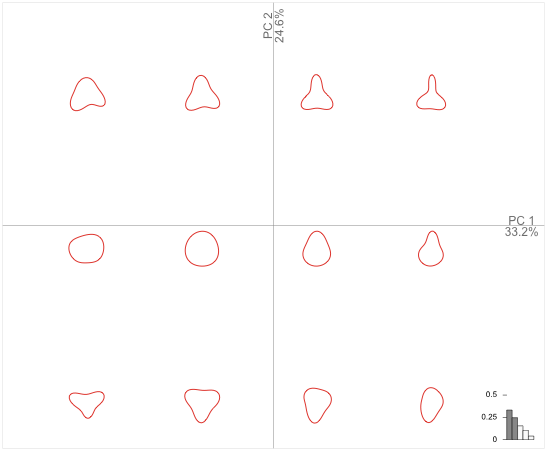

**Supp Fig. 3.** Shape variation derived from PCA on Elliptical Fourier Analysis.

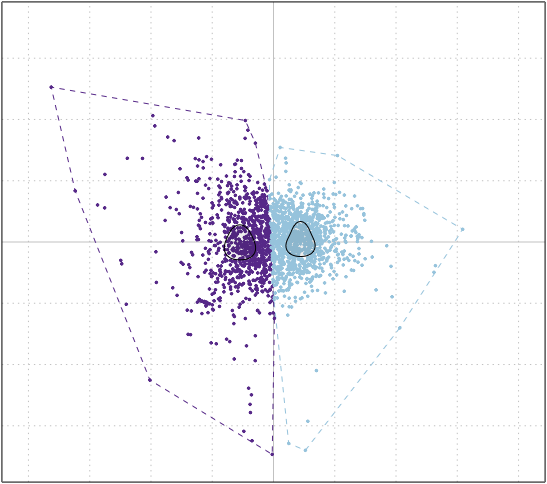

**Supp Fig. 4.** Clustering on the Elliptical Fourier components characterizing fruit shapes. When cluster number is set to two, the two characteristics shape are ‘elongated’ and ‘globose’ like, as shown by the black contours.

**Variational Autoencoder to describe shape categories:** We discovered shape categories from the latent space generated by a variational autoencoder deep neural network. The details are in the main manuscript. Here, we present the silhouette score of the clusters on the latent space varying between 2 to 9, the k-means clustering and two representative fruits of each cluster.

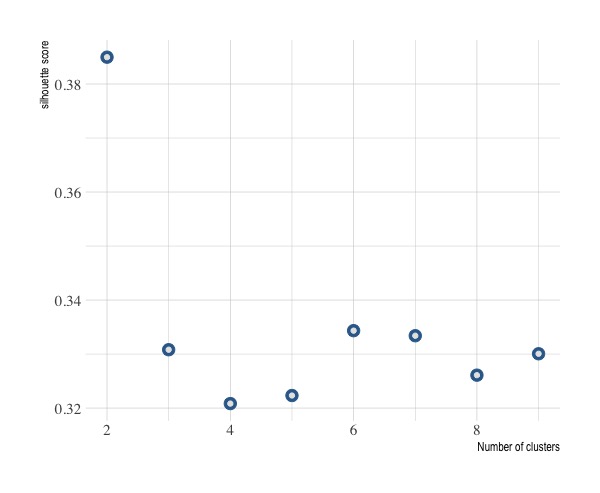

**Supp Fig. 5** Silhouette analysis for number of clusters on shape latent space from Variational autoencoders output. The index is the mean silhouette score through all the clusters. The optimal number of clusters is 2, i.e. there are two main shapes for strawberries in this database.

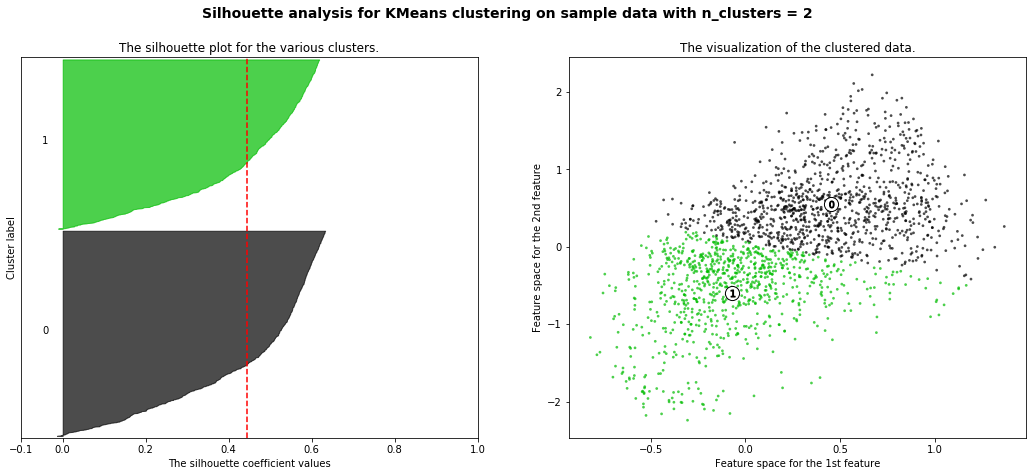

**Supp Fig. 6** Left: silhouette plot for each cluster (0 and 1). Right: visualization of the clustered data in the latent space.

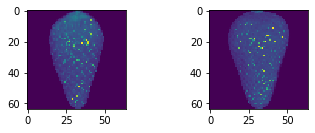

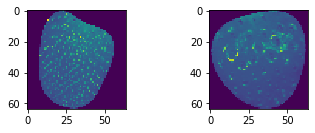

**Supp Fig. 7** The upper panel shows two examples of fruit belonging to the cluster 0, i.e. these are ‘elongated’ fruits. The lower panel show two examples of fruit on the cluster 1, these fruits have a ‘globose’ appearance.

Heritability evaluation

| **Trait** | **h2a** | **h2a_sd** | **h2d** | **h2d_sd** |
| --- | --- | --- | --- | --- |
| **Like- red** | 0.21 | 0.08 | 0.23 | 0.09 |
| **Pale** | 0.30 | 0.10 | 0.24 | 0.10 |
| **Like orange** | 0.18 | 0.07 | 0.20 | 0.07 |
| **Shape cluster** | 0.21 | 0.07 | 0.25 | 0.09 |
| **A_channel** | 0.20 | 0.07 | 0.20 | 0.08 |
| **B_channel** | 0.20 | 0.08 | 0.22 | 0.08 |
| **L_channel** | 0.20 | 0.07 | 0.26 | 0.09 |
| **Fruit height** | 0.22 | 0.08 | 0.23 | 0.09 |
| **Fruit width** | 0.16 | 0.06 | 0.24 | 0.10 |
| **Widht_at_75 height** | 0.18 | 0.06 | 0.25 | 0.10 |
| **Widht_at_25 height** | 0.16 | 0.05 | 0.21 | 0.10 |
| **widht_at_half_height** | 0.16 | 0.06 | 0.23 | 0.09 |
| **Area** | 0.16 | 0.06 | 0.25 | 0.10 |
| **Perimeter** | 0.21 | 0.07 | 0.20 | 0.08 |
| **Solidity** | 0.20 | 0.08 | 0.20 | 0.08 |
| **circularity** | 0.25 | 0.09 | 0.20 | 0.08 |
| **EllipseRatio** | 0.28 | 0.11 | 0.19 | 0.07 |
| **Hight/ width** | 0.28 | 0.10 | 0.22 | 0.09 |
| **Tip** | 0.21 | 0.08 | 0.24 | 0.09 |
| **Neck** | 0.21 | 0.08 | 0.20 | 0.07 |
| **Left side** | 0.21 | 0.08 | 0.22 | 0.08 |
| **Right side** | 0.21 | 0.08 | 0.26 | 0.10 |
| **Elliptical Fourier PC1** | 0.25 | 0.09 | 0.20 | 0.08 |
| **Elliptical Fourier PC2** | 0.25 | 0.09 | 0.27 | 0.10 |
| **Suppl. Table 2** Heritability values for all the shape and color related traits, the additive and dominant components were evaluated. | | | | |
